# Study of the molecular and spatial structure of trans-cinnamic acid isolated from the *hairy root* culture of *Scutellaria baicalensis*

**DOI:** 10.1101/2025.03.09.642299

**Authors:** Anastasia Mikhailovna Fedorova, Irina Sergeevna Milentyeva, Lyudmila Konstantinovna Asyakina, Alexander Yuryevich Prosekov

## Abstract

In this study, the crystalline structure of a sample of trans-cinnamic acid, isolated from an aqueous-alcohol extract of *Scutellaria baicalensis in vitro* root culture was analyzed using diffraction analysis. The diffraction analysis results confirmed that the investigated trans-cinnamic acid sample corresponds to the declared compound: trans-cinnamic acid. Additionally, the structure of this sample was examined using NMR spectroscopy. The obtained ^1^H, ^13^C{1H}, ^1^H^1^H-COSY, ^1^H^13^C-HSQC, ^1^H^13^C-HMBC NMR spectra of the studied sample correspond to the structure of the target compound, trans-cinnamic acid. No impurity signals were detected in the analyzed trans-cinnamic acid sample.

## Introduction

In recent years, the number of studies dedicated to identifying the key components in medicinal plant raw materials and their derived pharmaceutical preparations has been increasing. This trend is driven by the presence of phenolic compounds in these plants, which exhibit a wide range of biological activities widely utilized in medicine [1]. *Scutellaria baicalensis* (family *Lamiaceae*) is considered a valuable medicinal plant with a broad spectrum of applications in both traditional and official medicine. In scientific medicine, it is used as an antihypertensive and sedative agent for the treatment of hypertension (stages 1 and 2), as well as for functional disorders of the nervous system. *Scutellaria baicalensis* is known for its general tonic effects, its ability to stimulate the central nervous system, enhance the body’s resistance, regulate metabolism, and slow down the aging process [2]. The pharmacological effects of this plant are attributed to a wide range of biologically active compounds, including coumarins, tannins, essential oils, flavonoids, and many others. Special attention should be given to the group of phenolic compounds, due to their exceptionally high content and significant structural diversity [3–5].

With advancements in analytical techniques, the range of detectable compounds has significantly expanded, improving the reliability of biologically active substance (BAS) identification in medicinal plant raw materials. The most commonly used methods for this purpose include liquid chromatography (LC) combined with various detection techniques such as ultraviolet (UV) spectroscopy, infrared (IR) spectroscopy, nuclear magnetic resonance (NMR) spectroscopy, and mass spectrometry (MS). However, the number of studies utilizing these methods remains limited, and the available data are often inconsistent and contradictory [6–9].

It has been established that over the past two decades, NMR has become one of the three primary analytical methods, alongside gas chromatography (GC) combined with mass spectrometry (MS) and liquid chromatography (LC) combined with single-stage MS. Consequently, evaluating the structure of plant metabolites using NMR-MS is highly relevant [10], as LC methods are frequently used for the analysis of polyphenols.

The advantages of NMR-MS include high reproducibility, automation, fewer sample preparation steps, and built-in quantification of target compounds without the need for specific standards [11]. NMR spectroscopy allows for the detection of compounds that are not suitable for identification by LC-MS, such as various carbohydrates, organic acids, alcohols, polyols, and other highly polar compounds. The limitations and advantages of NMR are presented in Table 1 [12].

**Table 1.** Some key advantages and limitations of NMR compared to MS (GC-MS, LC-MS) [12].

| Criteria | NMR | Mass Spectrometry |
| --- | --- | --- |
| Reproducibility | High reproducibility | Lower reproducibility |
| Sensitivity | Lower sensitivity | High sensitivity |
| Selectivity | Usually used for non-selective analysis | In combination with LC, it is a good tool for targeted analysis |
| Sample measurement | Provides relatively fast measurement using 1D <sup>1</sup>H-NMR spectroscopy | To maximize the number of detected metabolites, various ionization methods are required |
| Sample preparation | Involves minimal sample preparation, usually transferring the sample into an NMR tube and adding a deuterated solvent. Can be automated. | Complex sample preparation, requires chromatography, derivatization is needed for GC-MS |
| Restoration example | Non-destructive method | Destructive technique |
| Quantitative analysis | + | MS signal intensity often does not correlate with substance concentration |
| Number of detectable substances | Detection and identification of fewer than 200 substances | Detection of thousands of substances and identification of hundreds |
| Targeted analysis | Possible, but usually used for non-targeted analysis | + |

One of the methods for studying the structure of substances is diffraction analysis, which includes X-ray crystallography and X-ray phase analysis. These methods are based on the diffraction of X-rays on a substance’s crystal lattice. X-ray crystallography is used for the analysis of single crystals, while X-ray phase analysis is applied to polycrystalline samples (powders). X-ray phase analysis determines which crystalline phases are present in the studied samples. The identification methodology involves detecting phases whose reflections correspond to those of the tested material. Qualitative X-ray analysis compares experimental diffraction data such as diffraction angles (2θ), interplanar spacings (d_hkl_), and relative intensities with values from reference databases [13].

The aim of this study is to investigate the molecular and spatial structure of trans-cinnamic acid, isolated from the hairy root culture of *Scutellaria baicalensis*, using NMR spectroscopy and X-ray phase analysis.

### Objects and methods of research

The object of this study is trans-cinnamic acid, obtained from the aqueous-alcoholic extract of the in vitro root culture of *Scutellaria baicalensis*.

Trans-cinnamic acid, isolated from the extract of *Scutellaria baicalensis* root culture, was obtained at earlier stages. The methods for cultivating the root culture of *Scutellaria baicalensis*, extraction, isolation, and purification of trans-cinnamic acid are described in the work of A. M. Fedorova and colleagues [14].

For the transformation of seedlings, grown for 14–28 days on a nutrient medium with the following composition: macroelements B5 – 50.00 mg, microelements B5 – 10.00 mg, Fe-EDTA – 5.00 ml; thiamine – 10.00 mg; pyridoxine – 1.00 mg; nicotinic acid – 1.00 mg; sucrose – 30.00 g; inositol – 100.00 mg; 6-Benzylaminopurine – 0.05 mg; indole-3-acetic acid – 1.00 mg; agar – 20.00 g. Also soil bacterial strains of agrobacterium rhizogenes 15834 Swiss (Moscow, Russia) were used. For the in vitro root cultures of Scutellaria baicalensis, the cultivation cycle lasted 5 weeks. The parameters for obtaining the aqueous-alcoholic extract of Scutellaria baicalensis root culture were as follows: 30 ± 0.2% ethanol, raw material-to-extractant ratio: 1:86, temperature: 70 ± 0.1°C, extraction time: 6 ± 0.1 hours. The scheme for the isolation and purification of trans-cinnamic acid obtained from the *Scutellaria baicalensis* root culture extract is presented in Figure 1. The purity level of trans-cinnamic acid was no less than 95%.

**Figure 1.**
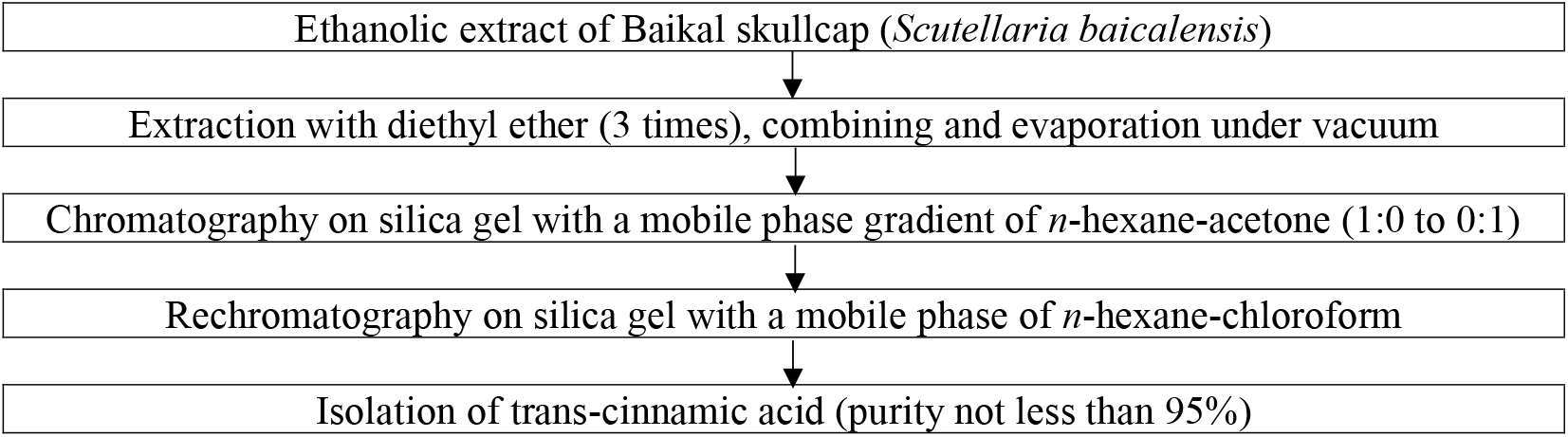
Purification scheme of trans-cinnamic acid obtained from the root culture extract of *Scutellaria baicalensis*

For the analysis of the crystalline structure of the research objects, an X-ray diffractometer Bruker D8 Advance (copper radiation CuKα) with a LynxEye position-sensitive detector was used. The measurement was conducted in reflection geometry with rotation. Data collection was carried out using the Bruker DIFFRACplus software package, while the analysis was performed using EVA and Topas V5.0 software.

For NMR spectroscopy, 20 mg of trans-cinnamic acid sample was dissolved in 600 µL of dimethyl sulfoxide (DMSO) at a temperature of 25°C. The resulting solutions were placed in a Bruker Avance IIIHD 500 NMR spectrometer for NMR spectrum registration. The operating frequencies of the NMR spectrometer are shown in Table 2.

**Table 2.**
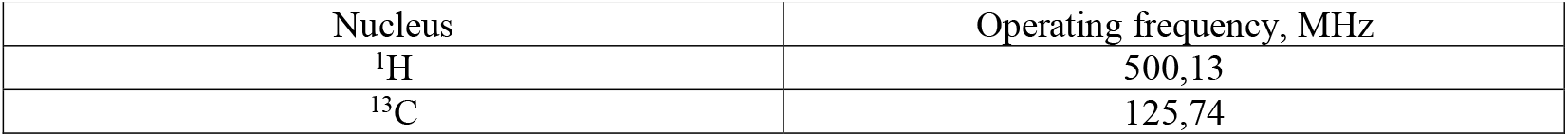
Operating frequencies used in the study.

The ^13^C{^1^H} NMR spectra were recorded in a decoupling mode to suppress spin-spin interactions with ^1^H nuclei, using phase selection (jmod-echo) to distinguish CH, CH_3_ / CH_2_, and quaternary C groups in the spectra.

To assign signals in the ^1^H and ^13^C{^1^H} spectra, additional 2D NMR spectra were recorded, including ^1^H^1^H-COSY, ^1^H^13^C-HSQC, and ^1^H^13^C-HMBC.

## Results and discussion

The analysis results showed that the trans-cinnamic acid sample is crystalline and single-phase, as presented in Figure 2 and Table 3.

**Table 3.**
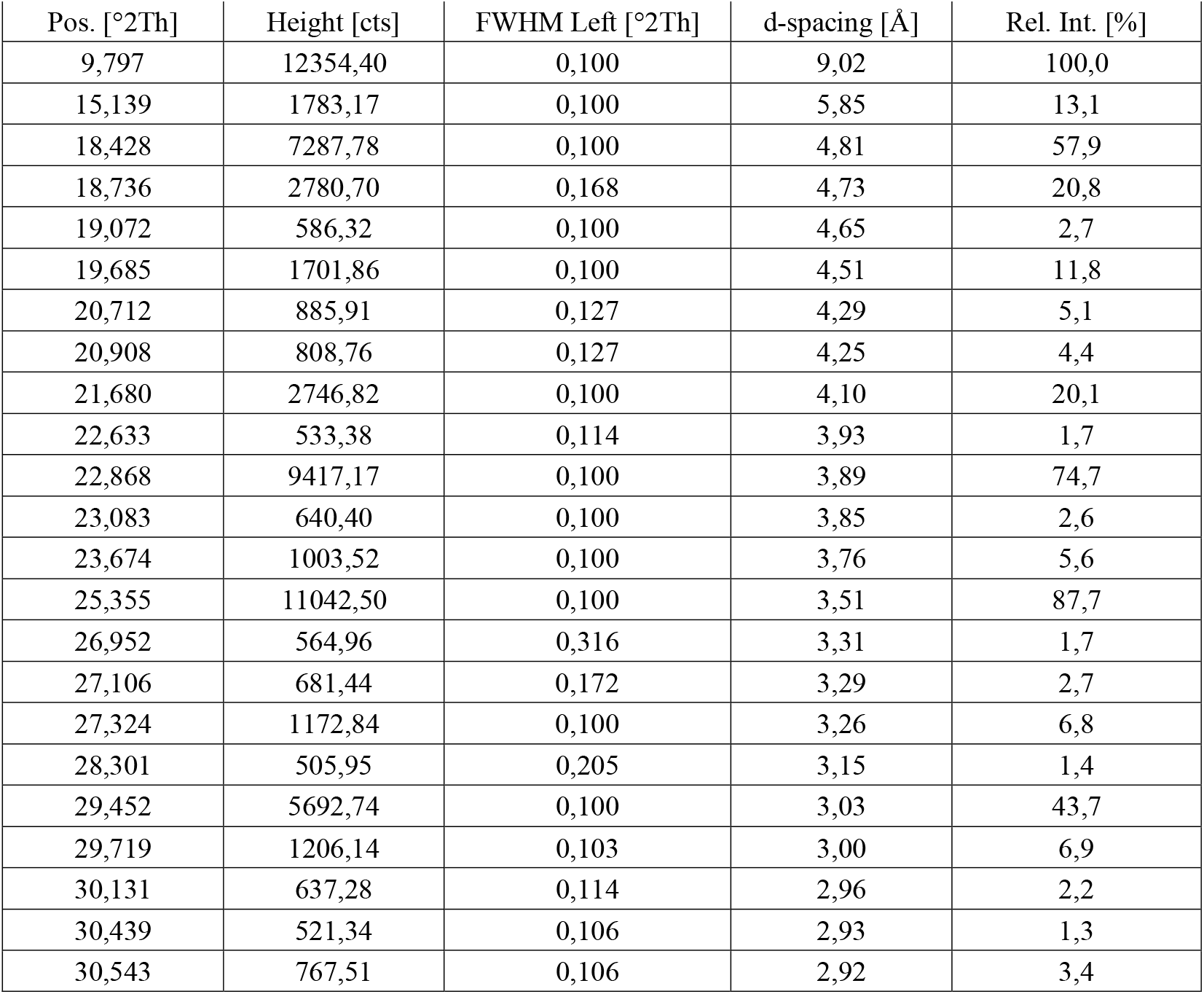
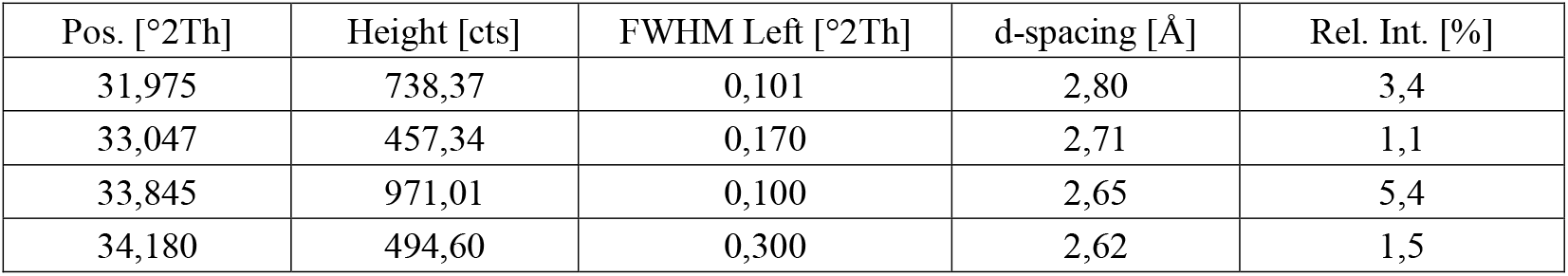
Intensity and position of selected peaks with a relative intensity greater than 1% for the trans-cinnamic acid sample.

**Figure 2.**
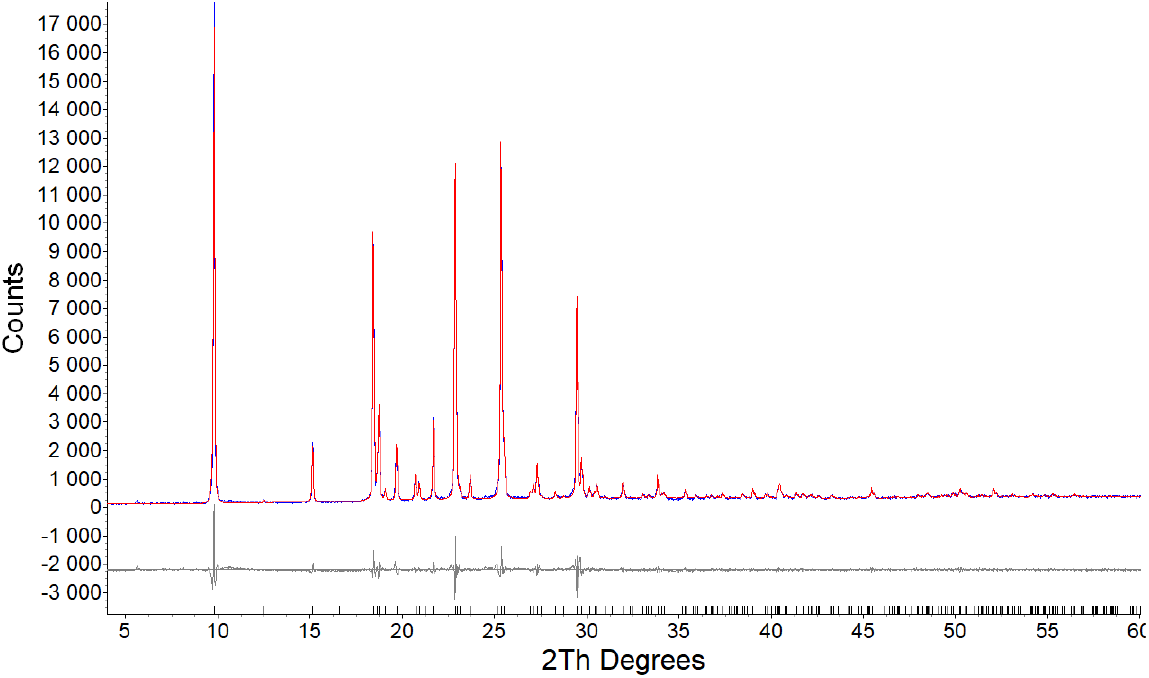
Experimental diffraction pattern of the trans-cinnamic acid sample (blue curve) and the theoretical diffraction pattern calculated for its unit cell (red line), along with their difference curve (gray line). Vertical ticks indicate the calculated peak positions.

The trans-cinnamic acid sample corresponds to the monoclinic phase of cinnamic acid (refcode in CCDC: CINMAC). Refinement results using the Pawley method: space group P2_1_/c, a = 5.6383(2) Å, b = 18.0163(4) Å, c = 9.0081(4) Å, β = 121.461(3)°, V = 780.53(5) Å^3^, R-Bragg = 0.623%, R_exp_ = 4.53%, R_wp_ = 9.20%, R_p_ = 6.78%, GOF = 2.03%.

The results of the diffraction analysis confirmed that the studied trans-cinnamic acid sample corresponds to the declared substance: trans-cinnamic acid.

Further structural investigations of the trans-cinnamic acid sample were carried out using NMR spectroscopy.

Figure 3 presents the NMR spectra of the trans-cinnamic acid sample, presumably having the structure of trans-cinnamic acid (^1^H, ^13^C{^1^H}, 2D ^1^H^1^H-COSY, ^1^H^13^C-HSQC, ^1^H^13^C-HMBC).

**Figure 3.**
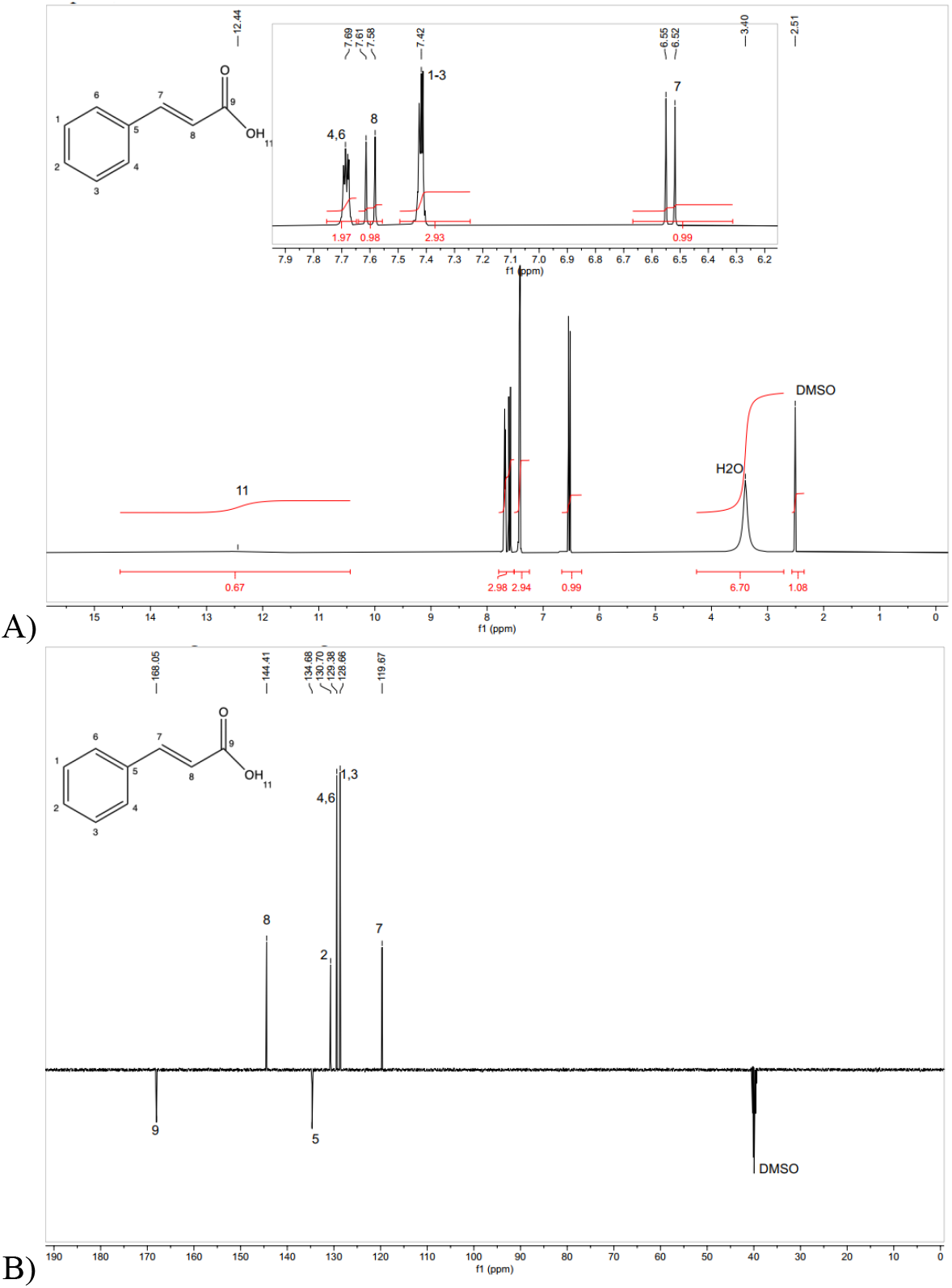
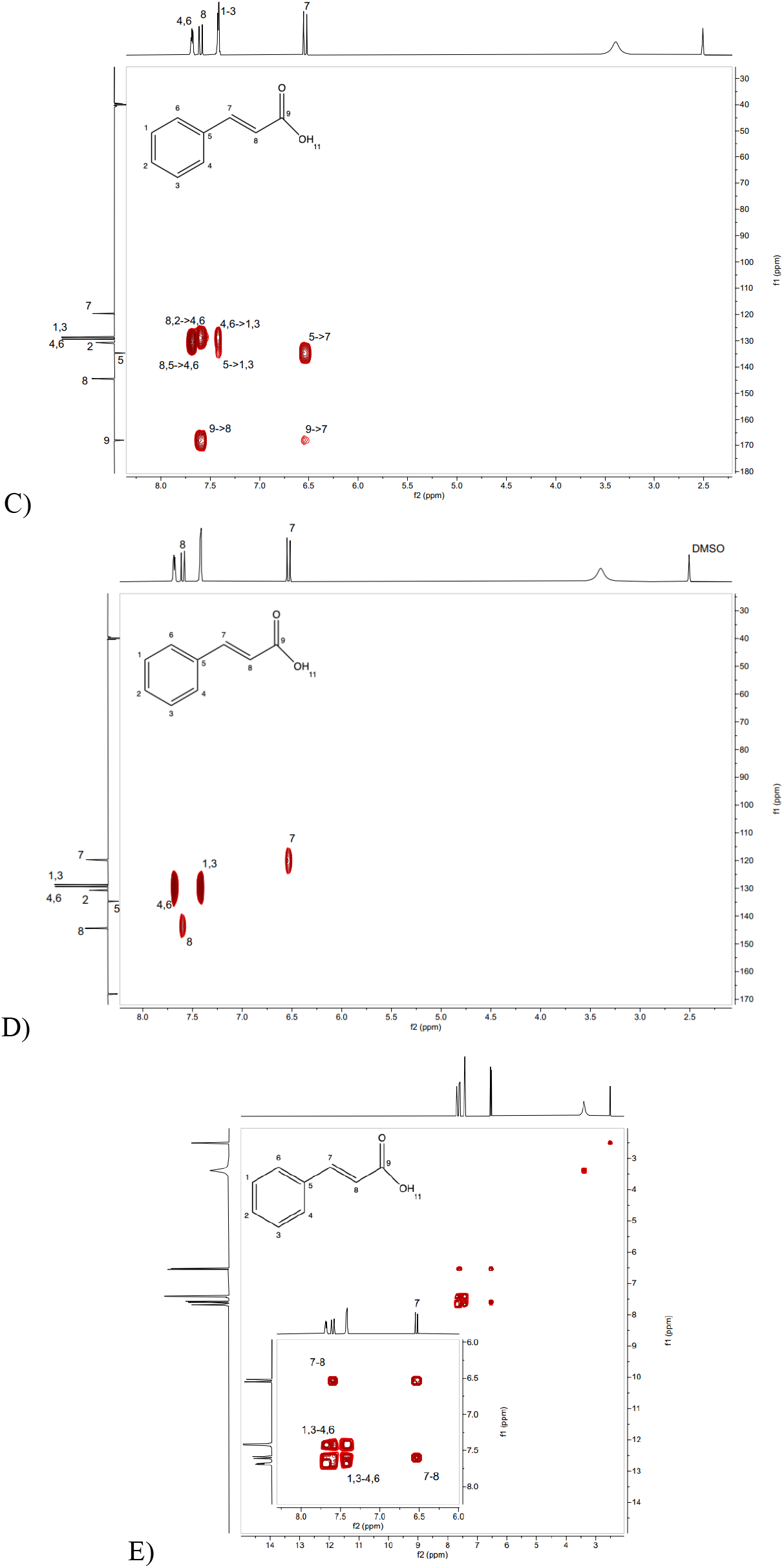
NMR spectra of the trans-cinnamic acid sample: A –^1^H; B –^13^C{^1^H}; C –^1^H^13^C-HMBC; D –^1^H^13^C-HSQC; E – 2D ^1^H^1^H-COSY

From the figure above, it is evident that the NMR spectra ^1^H, ^13^C{^1^H}, ^1^H^13^C-HMBC, ^1^H^13^C-HSQC, ^1^H^1^H-COSY of the presented trans-cinnamic acid sample correspond to the target product – trans-cinnamic acid, as shown in Figure 4. No impurity signals are present.

**Figure 4.**
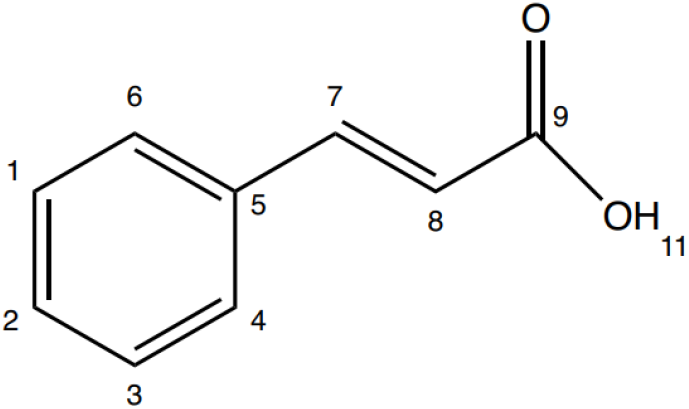
Structural formula of the trans-cinnamic acid sample identified as trans-cinnamic acid

Based on the number of signals, their positions in the spectra (chemical shifts), integral intensities, signal ratios, signal multiplet structures, and spin-spin coupling constants (SSCC), the obtained spectra correspond to the target product. The signal assignments are presented in Table 4 (^1^H and ^13^C{^1^H} NMR spectra).

**Table 4.** Signal assignments for the trans-cinnamic acid sample in the ^1^H and ^13^C{^1^H} NMR spectra.

| In the $^1\text{H}$ NMR spectrum | | |
| --- | --- | --- |
| $\delta$, ppm | xH, multiplicity, j (Hz) | Assignment |
| 6,53 | 1H, d, j=14.6Hz | CH(7) |
| 7,42 | 3H, m | CH(1), CH(2), CH(3) |
| 7,60 | 1H, d, j=14.6Hz | CH(8) |
| 7,69 | 2H, m | CH(4), CH(6) |
| 12,44 | 1H, bs | OH(11) |
| 119,67 | 1C, s | CH(7) |
| 128,66 | 2C, s | CH(1), CH(3) |
| 129,38 | 2C, s | CH(4), CH(6) |
| 130,70 | 1C, s | CH(2) |
| 134,68 | 1C, s | C(5) |
| 144,41 | 1C, s | CH(8) |
| 168,05 | 1C, s | C=O(9) |

The results of the conducted NMR spectroscopy confirmed that the studied object, trans-cinnamic acid isolated from the aqueous-alcoholic extract of the in vitro root culture of *Scutellaria baicalensis*, corresponds to the declared compound: trans-cinnamic acid. No impurities were detected in the trans-cinnamic acid sample.

## Conclusion

This study aimed to assess the molecular and spatial structure of trans-cinnamic acid isolated from the extract of *hairy roots Scutellaria baicalensis* using NMR spectroscopy and X-ray phase analysis. The experiment established that the NMR spectra (^1^H, ^13^C{^1^H}, ^1^H^1^H-COSY, ^1^H^13^C-HSQC, ^1^H^13^C-HMBC) of the studied sample correspond to the structure of the target compound – trans-cinnamic acid. No impurity signals were detected in the trans-cinnamic acid sample.

## Funding

*This work was carried out as part of the state-funded project «Development of biologically active supplements consisting of metabolites from plant in vitro cultures to protect the population from premature aging» (project FZSR-2024-0008)*.

*The study was conducted using the equipment of the Shared Research Facility «Instrumental methods of analysis in applied biotechnology» at Kemerovo State University*.

